# Determination of secretion type associated motifs from pathogenic protein sequences

**DOI:** 10.1101/2022.06.10.495625

**Authors:** Orhan Özcan, Gökalp Çelik, Seren Sert, Gülay Özcengiz, O. Uğur Sezerman

## Abstract

Currently, potential vaccine candidates are determined using protein localization predictors to identify surface or secreted protein sequences. The pathogenic bacterial sequences constitute only a tiny percentage of this database. They search for general motifs found in bacteria and miss out on pathogen-specific motifs, which is crucial for identifying vaccine candidates. We developed a motif search algorithm derived from pathogenic bacterial sequences in this work. Using this method, we identified secretion motifs of pathogenic bacteria for six different secretion pathways and non-specific secretion motifs. To this end, we established a database of immunogenic sequences from patented vaccine sequences, AntigenDb, and PUB-MED. The motifs are filtered out according to their presence in a cytoplasmic dataset to keep motifs that are present only in secreted sequences. These sequences are validated on newly identified novel vaccine candidates from 20 pathogenic bacterial immunoproteomics data.

## 1. INTRODUCTION

The subcellular localization of a protein provides essential insights into the function of a protein [1]. There are several wet-lab methods like green fluorescent protein (GFP) tagging, which gives detailed information on the localization of proteins; however, when massive proteome data is considered, high throughput screening methods are needed to analyze the data in a fast, automated, cost-efficient manner with high accuracy. More than 40 prediction tools have been used for protein localization prediction. Nonetheless, most bacterial localization prediction (BLP) tools end up with false-positive results in exchange for better sensitivity [2]. Although most localization tools can successfully execute the prediction, especially for non-pathogenic bacteria, by using the Conserved Domain Database (CDDs) for localization prediction, there may be some misleading results for pathogenicity-related proteins due to using CDDs. For example, bacteria send effector proteins to bind the ribosome, intervening in the host’s protein synthesis as part of their pathogenicity. Since it has a ribosome binding domain, the prediction algorithms classify it as a cytoplasmic protein. Bacteria secrete the protein as part of its pathogenicity. Even highly studied Diphtheria Toxin gi|157833808 is classified as a cytoplasmic protein by cellular localization predictors because of its action mechanism held directly inside the host cytoplasm.

Among the considerable variation of all bacterial species, pathogenic bacteria only constitute a small portion of bacterial sequence space; hence, it becomes statistically unimportant to include these sequences for overall prediction accuracy. PSORT program is the first prediction tool designed and is still one of the foremost tools used in bioinformatics; however, for the same reasons mentioned above, this tool can give fallacious results for pathogenic sequences.

This study shows a PSMS algorithm we have developed for identifying motifs in pathogenic sequences. As such, we could determine whether potential vaccine candidates were mistakenly predicted as cytoplasmic by other predictors or excluded from vaccine research. PSMS program is now available and can be used for scientific research in immunoproteomics, reverse vaccinology, and other molecular science areas for pathogenic bacteria. We created a database of pathogenic sequences by extracting all of the sequences from AntigenDb [3] and including other patented vaccine sequences. As patented sequences are highly focused databases, their usage in algorithms protects several pathogenic sequences from being diluted by localization predictors.

We have also tested PSMS with Bordetella pertussis sub-proteome data of our research group’s

## 2. MATERIALS AND METHODS

### 2.1 Database Construction

This study used AntigenDb, PUBMED, and USPTO to construct pathogenic datasets. Four hundred eighty-one prokaryotic pathogenic sequences were obtained from AntigenDb. We searched the patented vaccine sequences from USPTO and gathered 5166 additional pathogenic subproteome sequences. Proteins whose sequence homology is higher than 50% with the other sequences in the database were filtered using the CD-HIT program [5], resulting in 1740 additional pathogenic proteins to the AntigenDb database. Secretion pathway information was retrieved for each pathogenic sequence using Geneious R9 [6]. Based on their secretion type, we divided our data into six different datasets, namely Type 1 secretion associated protein dataset (T1S), Type 2 secretion associated protein dataset (T2S), Type 3 secretion associated protein dataset (T3S), Type 4 secretion associated protein dataset (T4S), Type 5 secretion associated protein dataset (T5S), and Type 6 secretion associated protein dataset (T6S).

Another database of 13308 truly cytoplasmic proteins that do not have immunogenic activity has been established to verify that it can genuinely distinguish secreted proteins. The number decreased to 2582 soon after the removal of homologous proteins by CD-HIT. Then to test the prediction ability of our algorithm in an independent data set, we have gathered 70 immunogenic proteins of 20 common human pathogens from recent immunoproteomic papers in PUBMED. After the removal of homologous proteins by CD-HIT, the number has decreased to 66. The pairwise identity of these proteins was found to be 6% by Multiple Alignment with only 0.3% identical functional sites, which states that these proteins are evolutionary distant pathogenic proteins. A distant pathogenic protein set is desired for program efficiency tests. Both the PSMS program and the databases used in this research are available for the scientific community as supplementary files.

#### 2.1.1 Sub-proteomic Data of our group

We had identified sub-proteomic from Bordetella pertussis, which has the highest non-cytoplasmic over cytoplasmic protein ratio with respect to several bacterial sub-proteomes [4]. However, cytoplasmic contamination is inevitable with the current proteomics applications. To develop precise localization patterns, one must accurately construct proteomics data.

“Spectral counts” and “relative abundance,” derived from the nano-LC-MS/MS analysis of peptides, are two significant components in data filtering. BP0490 DNA polymerase III subunit beta was selected as the reference protein because its concentration is the highest of all the cytoplasmic leakage proteins. Thus, protein concentrations higher than the BP0490 DNA polymerase III subunit beta were selected for pattern analysis. Through filtering proteins, more accurate surfacome and secretome data were obtained. Although this step is crucial to obtaining precise sub-proteome data, we also missed some rare surface proteins as a side-effect. Out of 210 initially defined proteins after the filtering step, we ended up with 45 secreted proteins. This Data is used as independent data to find non-specific secretion system patterns.

### 2.2. Pattern Development

PSMS is based on identifying motifs (conserved sequences) in protein families with similar secretion pathways. The optimum motif length is selected between 5 and 18 amino acids. Five long amino acids were selected because one well-known localization signal LPTXG has conservation of 5 amino acids. Eighteen amino acid motives are the most extended prokaryotic motifs derived from the work by Katti et al. [7]. There are also more extensive conserved bacterial motifs, but they are mainly membrane-bound proteins and their membrane-spanning repeats. While grouping these k-mers, the allowed number of similar groups of amino acids and the allowed number of a different group of amino acids in k-mers were derived from the Analysis of PSORTv3b motifs. PSORTv3b motifs are aligned for each maximum length number of different mutations, and the number of identical amino acids is determined from each multiple alignment. These numbers are given as parameters as tools for the PSMS algorithm.

The dataset was arranged for the five-fold test. Then for the train protein dataset (%80 of each dataset), all the k-mers sequences (ranging from 5-to 18) are obtained in a sliding window. In other words, the dataset is randomly divided into five groups, and the rules are determined from 4 groups and tested on the new 5th group.

Similar to all k-mer dependent pattern construction algorithms, our first output was more significant than the resource databases. For example, the 9-aminoacid length pattern has 8-, 7-, 6-, and 5-aminoacid sub-string versions. The pattern [IVALF]NXXGGXXG has 159 hits for all versions. The highest string sized of these versions is used for web-logo projection of the patterns so that RegEx of the patterns can be created. Web-logo projection categorizes amino acids into five different groups with respect to their physicochemical properties of the amino acids. If any position has three or more categories, we have used X as no specific physiochemical property is needed.

All the RegEx formed patterns that were filtered with the help of a cytoplasmic dataset. If a motif is present in any cytoplasmic dataset, it is filtered out. After this filtering, another filtering step is done for specificity. Suppose a motif is present in more than one type of secretion system dataset. In that case, that motif is classified as a non-specific secretion system motif or else stays as a specific type of secretion system motif. Concerning table 5 and table 6, 57 patterns were divided into six specific and non-specific secretion systems. Secretion system patterns that cannot satisfy specificity and those created by using the *Bordetella* surfacome dataset of our own are listed in the non-specific secretion system. Motives similar to each other under these constraints are considered secretion motif candidates. For a motif to be classified as a secretion motif, at least 5% of the proteins belong to the same secretion type dataset. As a report, the algorithm gives all the secretion types for any given sequence for all the secretion pathways.

#### 2.2.1 Pattern Development from our Data

The high ratio of non-cytoplasmic proteins in the previous study [4] of our research group led the team to examine LC-MS data concerning the spectral counts and relative abundances. After careful data filtering in 2.1.1, we did the secretion system-specific pattern construction process until RegEx development. Unlike the other datasets we created for the secretion systems, our data consist of only single bacterial proteins. In order to validate pattern candidates, valid for *Bordetella pertussis*, for other pathogens, we performed taxonomy BLAST.

[Table2, 3, 4 and Figure 2]. We have downloaded the hits for each taxonomy and used these sequences for the RegEx development of pattern candidates. Constructed ReGex was also filtered out by cytoplasmic dataset presence.

**Figure 1.**
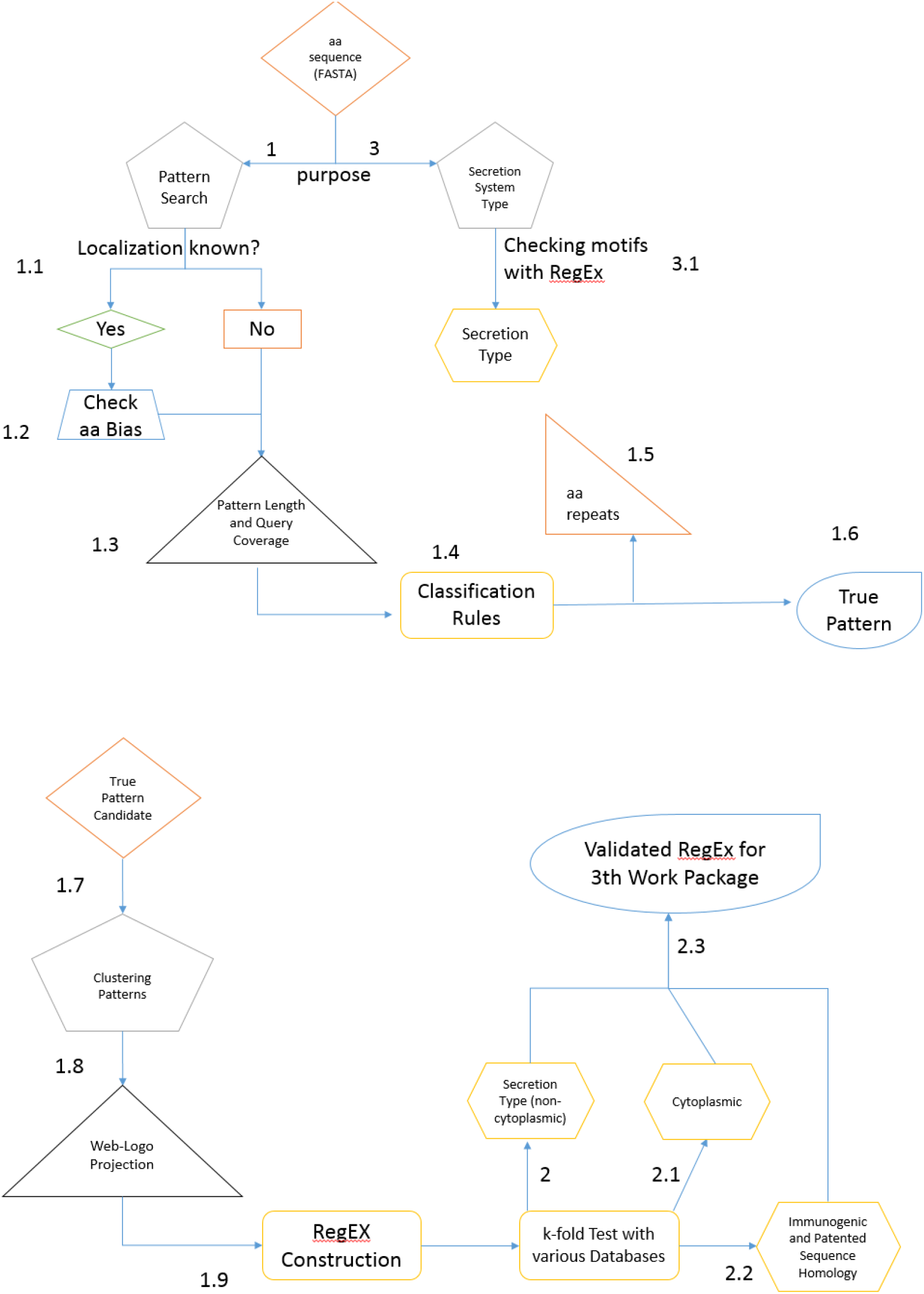
Flow-chart of Pattern development.

**Figure 2.**
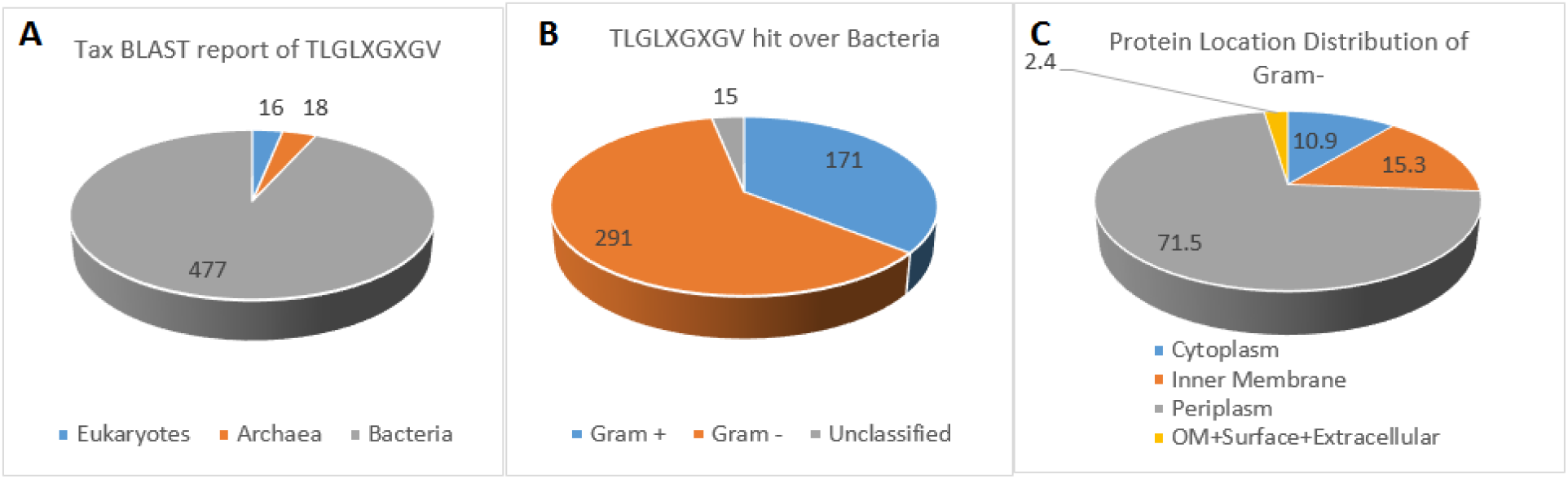
BLASTP analysis of the TLGLXGXGV pattern; A) Distribution of protein hits among Kingdoms, B) Distribution of proteins among bacteria, C) Distribution of Gram (-) protein hits with respect to cellular localization.

### 2.3. Repeat Analysis

Initially, we aimed to find a locational pattern from the sub-proteomic data. We figured out that several pattern candidates were not good patterns; instead, they were dipeptide and tripeptide repeats. We have implemented the “short pattern candidates” (SPCS) function to PSMS to remove dipeptide and tripeptide. The relation between protein functions and their localization concerning their short amino acid repeats was shown by several research groups [8, 9, 10, and 11].

## 3. RESULTS

### 3.1 Pattern Analysis of Bordetella pertussis

TLGLXGXGV, TXALAVAG, PXN and [VALST][ANILV][LAVM][GT][ATGVPL][NTSG][ALNST]X[AVIW][TSPM] patterns are found in our Bordetella pertussis sub-proteome. Instead of having specific secretion associated dataset, we have a clear surfacome having a mixture of secretion associations, so we have focused mainly on cellular localization. If the triplets are LGX, RGX, and AGA for the TLGLXGXGV, the proteins are non-cytoplasmic regardless of classification. Moreover, for the TXALAVAG, if the mismatched amino acids are leucine, valine, and serine, the protein is non-cytoplasmic. Major unknown secretion patterns (hemagglutinin type, Hsp60, and EF-Tu) might be the products of exosomes. Our previous immunoproteome research group has shown their exosome-linked secretion [4]. The trojan horses of the pathogens, exosomes, are probably the essential elements regulating or impairing the toll-like systems.

### 3.2 Amino acid Repeat Analysis

In this study, the PNN repeat, previously shown in Katti et al. [7], was improved as PXN where X can be N, D, and S, but mostly D (Table 3 and Table 4). Functional domains carrying sequence abundance are in Table 4, showing that PXN tends to be present in binding domains. The PXN repeats are mainly found on the pathogens’ surface as virulence factors. Some o these repeat-bearing virulence factors are the peptidases and collagenases. Although PSORTv3b prediction results show these proteases as cytoplasmic proteins, these virulence factors are located on the pathogen’s surface or secreted.

### 3.3 Testing PSMS

While testing PSMS, the immunogenic protein dataset not used in pattern construction was used. Twenty pathogen genera with 70 immunogenic proteins are reduced to 66 proteins after application of the CD-HIT program to remove homologous sequences (%50 identities). Out of sixty-six pathogenicity-related proteins, only twenty-one are addressed for secretion systems. The other 45 sequences did not contain any secretions PSMS motifs that we searched for, so they were not classified as pathogenic. Twelve of the twenty-one proteins are addressed to a specific secretion system. Three of the remaining nine proteins had more than one PSMS motif, and the rest were matched with undetermined secretion systems. Type 3 and Type 4 secretion system association patterns were detected with PilQ of *Neisseria meningitis* and *Salmonella enterica*. Multiple secretion system usages by this protein are shown by researchers [12]. Another multiple secretion systems giving protein is PilT which shares similar routes with PilQ.

Immunogenic proteins such as LSU ribosomal protein L12P of *Chlamydia trachomatis*, protein translation elongation factor g of *Brucella melitensis*, cell envelope integrity inner membrane protein TolA of *Klebsiella pneumoniae*, chaperonin GroEL of *Legionella pneumophila*, 8-amino-7-oxononanoate synthase of *Mycobacterium tuberculosis*, elongation factor Tu of *Neisseria meningitides*, are classified as unknown secretion type proteins by PSMS. Unclassified secretion pattern is conserved in secreted proteins where they can not differentiate their secretion route.

Although we do not have a wet-lab proof for the unclassified secretion types, they are most probably the products of the exosomes. This type of cytoplasmic protein is found to be immunogenic in various studies, and its presence is also shown by various outer membrane vesicle proteomes of various pathogens. Based on the functional domains by the protein localization prediction programs, these proteins are classified as cytoplasmic, and many of them have been ruled out nearly two decades from wet-lab assays as vaccine candidates.

### 3.4 Result of Test Dataset on both PSMS and PSORTv3b

We built databases of pathogenic sequences [Table1] and searched for selected 5 to 18 amino acid long motifs as a new approach, namely Pathogenic Sequence Motif Search (PSMS). The algorithm found 56 distinct secretion-associated patterns and one three amino acid peptide repeat covering six different secretory pathways for predicting surface and secreted proteins. The cytoplasmic protein dataset, on the other hand, was used to exclude specific candidate patterns. Hundred and one patterns out of 158 pattern candidates, also observed in a cytoplasmic dataset, were excluded. Fourteen patterns from non-excluded secretion association patterns were regrouped for not satisfying secretion type specificity (Table 5& Table 6). These fourteen patterns are also crucial as they can be used for unknown secretion system proteins. We have found that these fourteen patterns are present in several outer membrane vesicle proteins of pathogens such as chaperonin, Elongation factor Tu and TolB proteins. Outer membrane vesicles are classified as a newer secretion system (Perez-Cruz et. al., 2013 [31]), and our findings confirm these observations. This new system is generally similar to the Type 3 secretion system. Instead of using microinjection by the pathogen to the host cell to change the hosts’ metabolism for improved pathogenesis, it uses outer membrane vesicles.

**Table 1.** Database created in this study.

| Database | # Of Proteins after CD-HIT filtered 0.5 |
| --- | --- |
| T1SS | 954 |
| T2SS | 668 |
| T3SS | 381 |
| T5SS | 770 |
| T6SS | 221 |
| Secreted (orphan secreted protein database) | 2533 |
| Total Secretion System Related | 5774 |
| Cytoplasmic | 2582 |
| 20 Common Human Pathogen Immunegic Proteins from PUBMED | 66 |

**Table 2.** BLASTp hit distribution of TLGLXGXGV pattern among prokaryotic organisms

| Kingdom | Class | # of Org. | Total Hit | Amino acids of <u>XGX</u> in Pattern |  |  |  |  |  |
| --- | --- | --- | --- | --- | --- | --- | --- | --- | --- |
|  |  |  |  | UN | C | IM | P | S,EC | Location |
| Archaea | *Euryarchaeotes*<br>-<br>*Crenarchaeotes* | 18 | 25 | 2 | 14 | 9 | - | - | AGF,VGG are inner membranous |
| Bacteria | CFB group | 28 | 37 | 1 | 4 | 32 | - | - | All inner membranous ones have LGY |
| Bacteria | γ-*proteobacteria* | 40 | 49 | 4 | 9* | 17 | 14 | - | LGL and RGG are membranous and periplasmic |
| Bacteria | α-*proteobacteria* | 21 | 25 | 4 | 6 | 12 | - | 3 | TLG inner membranous |
| Bacteria | β-*proteobacteria* | 181 | 241 | - | - | 12 | 229 | - | IGX,RGX are periplasmic |
| Bacteria | Δ-*proteobacteria* | 23 | 38 | 2 | 28 | 8 | - | - | TGL,VGG,VGC cytoplasmic |
| Bacteria | *Planctomyces* | 2 | 2 | - | - | 2 | - | - | Insufficient data |
| Bacteria | *Cyanobacteria* | 6 | 5 | 2 | 3 | 1 | - | - | Insufficient data |
| Bacteria | *Firmicutes* | 121 | 144 | 4 | 7 | 74 | 1 | 35 | NGD surface |
| Bacteria | *Deinococcus-Thermus* | 5 | 9 | - | 1 | 5 | - | 3 | LGX inner membranous |
| Bacteria | *Actinobacteria* | 47 | 62 | 2 | 15 | 40 | - | 5 | LGX inner membranous |
| Bacteria | Unclassified bacteria | 1 | 2 |  | - | 2 | - | - | Insufficient data |
| Total |  | 509 | 674 | 26 | 90 | 215 | 244 | 47 | LGX, RGX, AGA are non-cytoplasmic regardless of classification |

**Table 3.**
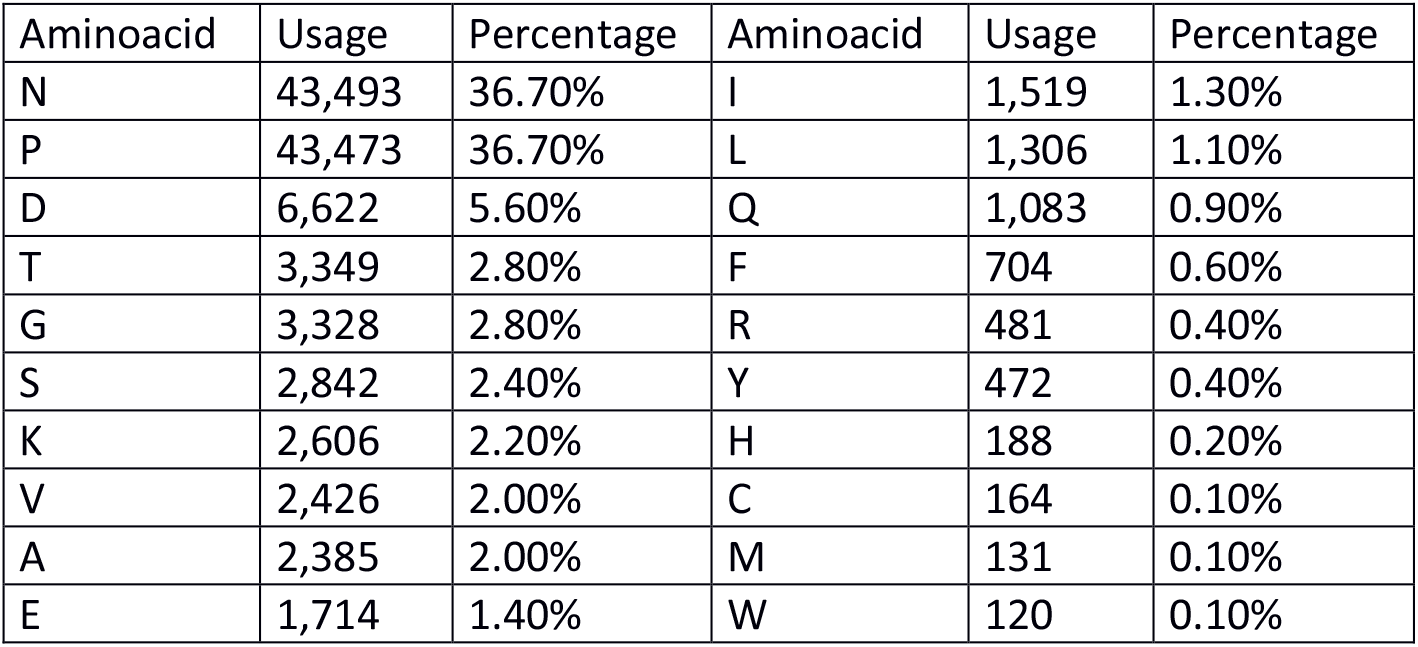
The amino acid usage frequency of PXN repeat in Bacterial PDB at NCBI.

**Table 4.**
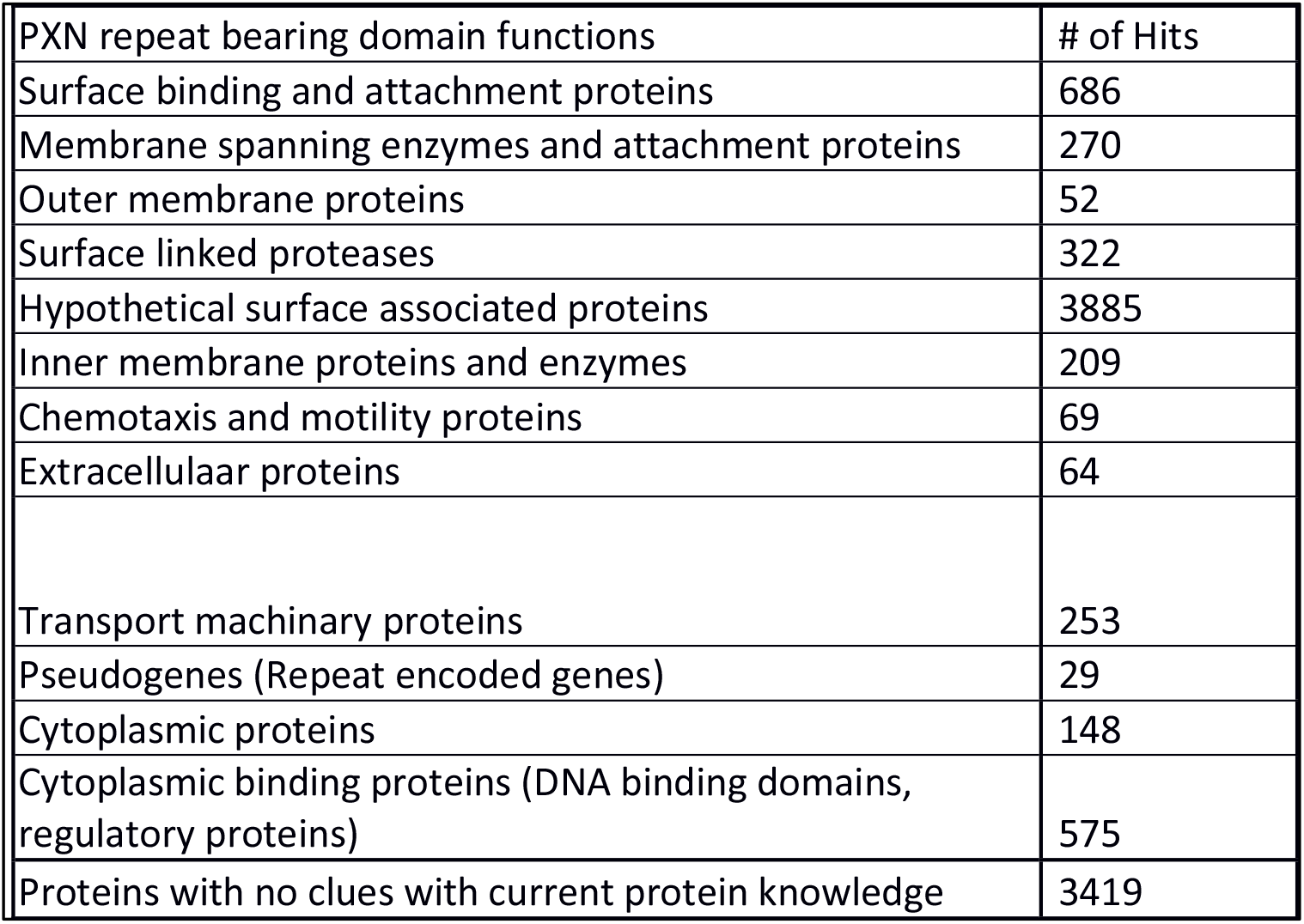
Functions of triple and quadruple PXN repeat bearing domains where BLASTp finds in NCBI non-redundant prokaryotic database.

**Table 5.** K-mers to Specific Patterns.

|  | Total # of Strings&Substring Hits | # Of Pattern Candidate Hits | Web-Logo Construction From Hits | 5 Fold Assay Passed | Cytoplasmic Filtered | Specificity |
| --- | --- | --- | --- | --- | --- | --- |
| T1SS | 14805 | 3224 | 174 | 41 | 7 | 5 |
| T2SS | 18490 | 3032 | 154 | 34 | 9 | 9 |
| T3SS | 5814 | 1302 | 105 | 27 | 8 | 7 |
| T4SS | 5145 | 1137 | 127 | 32 | 12 | 10 |
| T5SS | 8162 | 799 | 51 | 14 | 8 | 7 |
| T6SS | 5518 | 982 | 60 | 9 | 5 | 5 |
| Non Spesific |  |  |  |  |  | 14 |

**Table 6.**
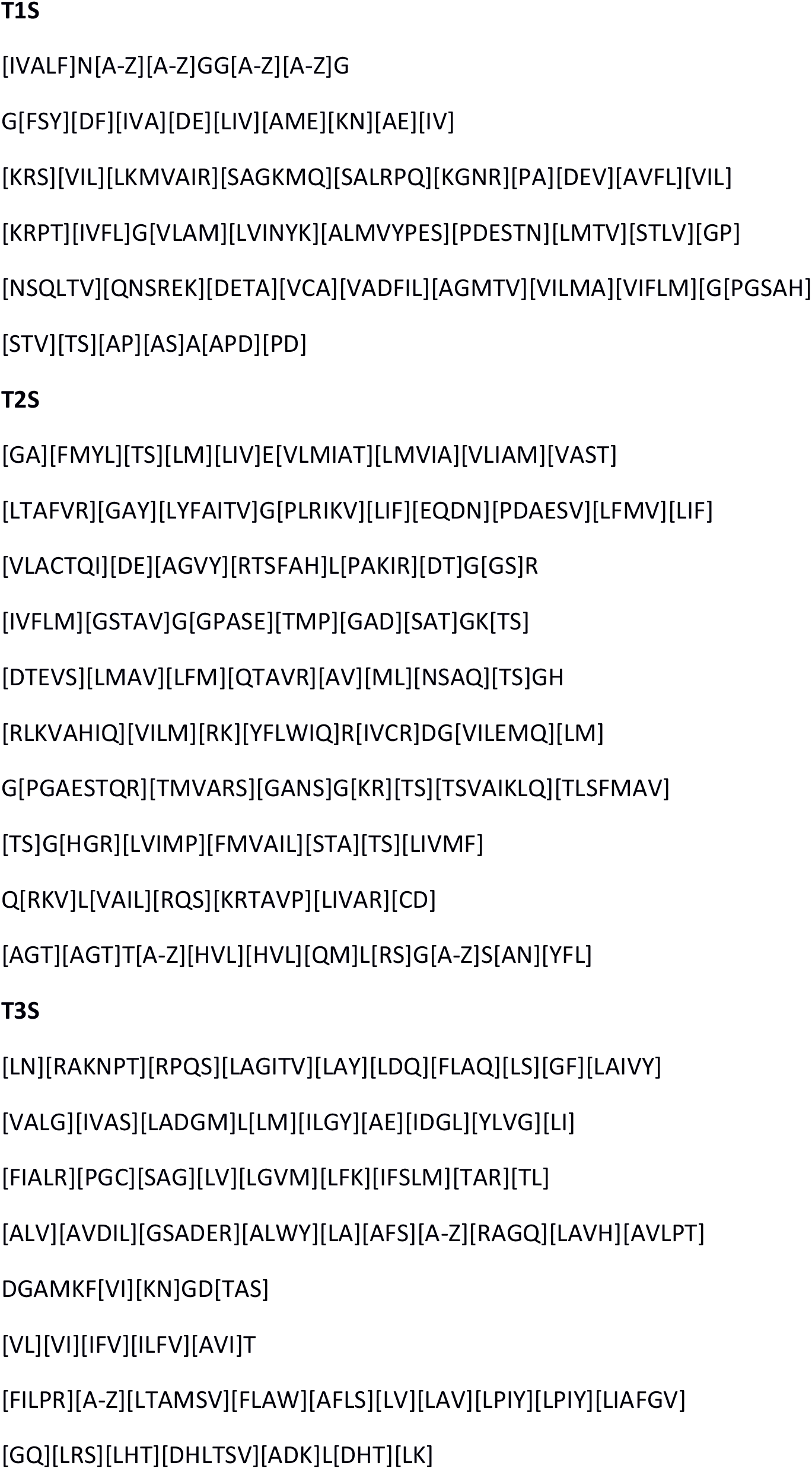
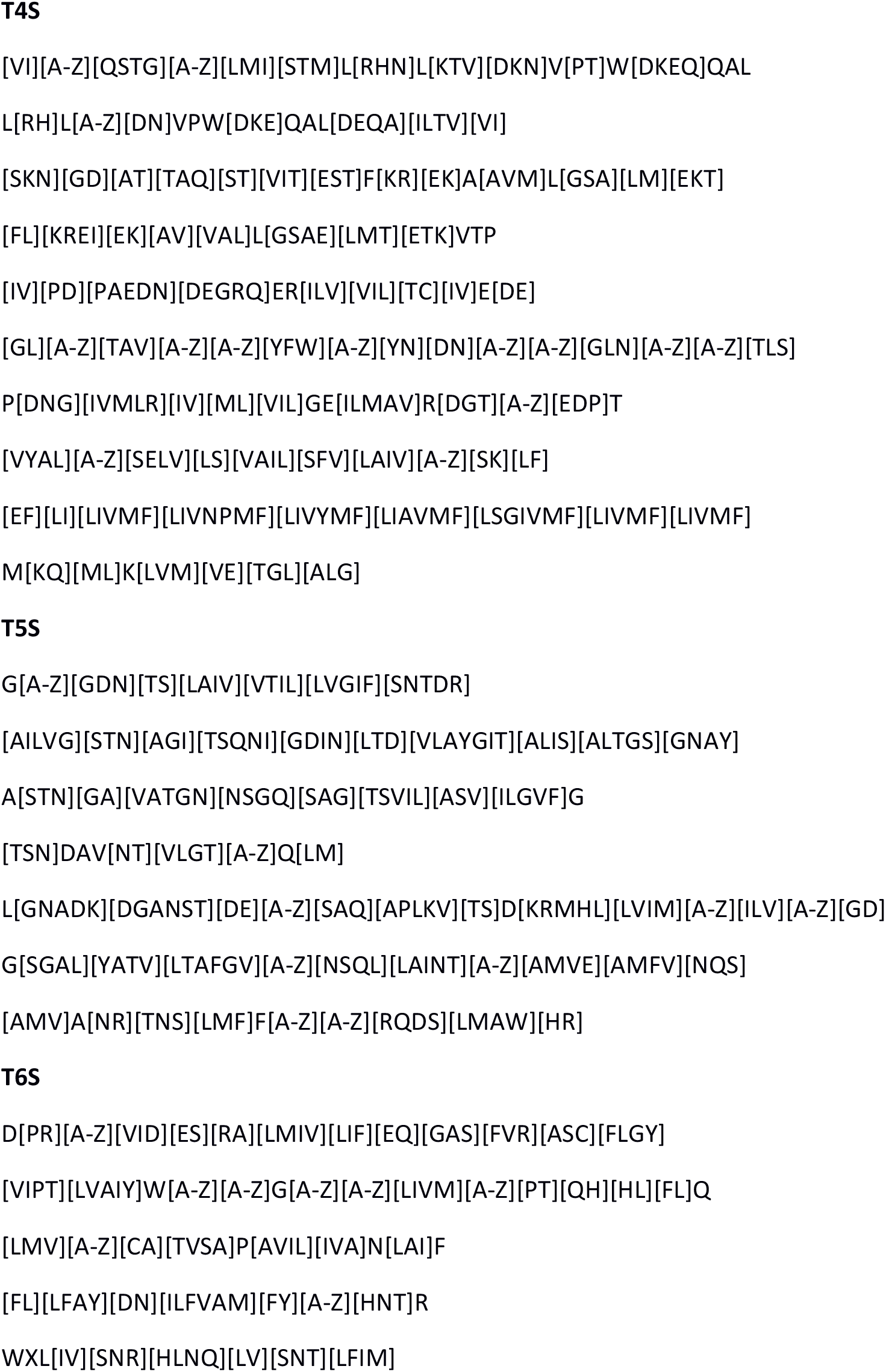
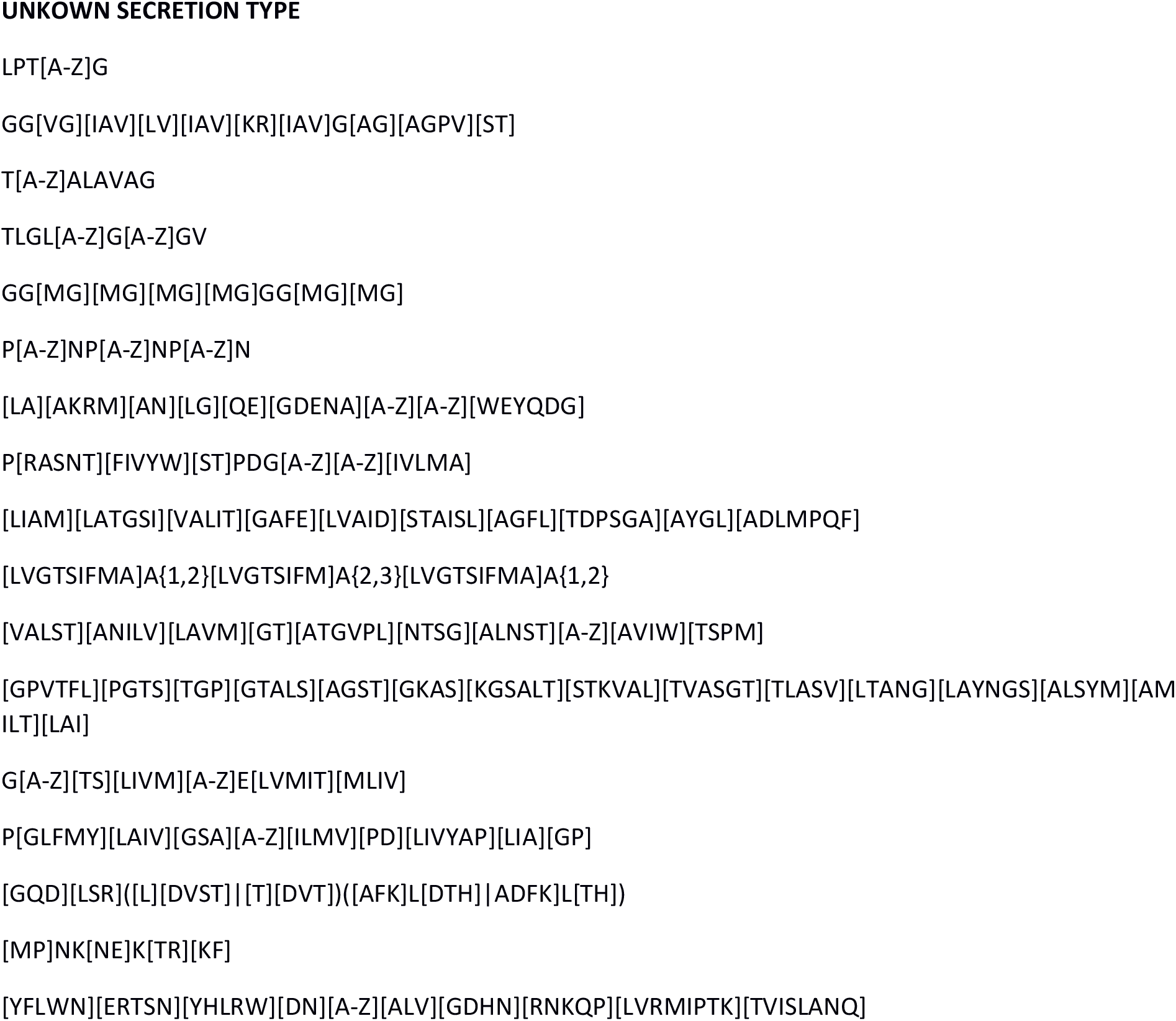
Patterns Found in this Research.

The immunogenic test dataset was run with PSORTv3b and PSMS. Eleven immunogenic proteins classified as unknown or cytoplasmic by PSORT are linked with secretion systems by PSMS. Six of the eleven proteins are linked with specific secretion types, and the rest are matched with an unclassified secretion system. The unclassified proteins are found mainly in various secretome studies and outer membrane vesicles studies. Instead of null functioning cytoplasmic proteins, various wet-lab experiments have discovered their pathogenicity-related roles. For example, GAPDH and enolase, classified as cytoplasmic proteins by localization predictors, have an affinity to binding cytoskeletal proteins of mammalian cells [13]. In the future, chaperonin GroEL, elongation factor Tu and FtsZ involved pathogenicity related secretion system might be clarified, and proteomic studies shall not explain their presence in secretome just because of cytoplasmic leakage.

Six proteins we could not show any relation with secretion in the test dataset were related to the secretion system by PSORT.

Both PSORT3b and PSMS could not respond to 13 proteins in the test dataset. The pathogenic test dataset can increase the overall correction of protein localization by 15% in the pathogenic test dataset.

## 4. DISCUSSION

Not only giving new patterns for pathogenicity-related secretion systems of bacteria, but also we are serving our particular protein databases to the scientific community. Moreover, the new secretion system might be enlightened from our non-specified secretion system pattern hitting proteins. With our patterns’ help, pathogenic proteins that were formerly predicted to have an intracellular localization and mistakenly ruled out as potential drug targets/vaccine candidates were successfully predicted as surface-associated/secreted. PSMS is the first tool to identify secretory motifs in secreted proteins and their secretion pathway. Besides, we have established extensive databases with respect to immunoreactivity and distinct secretory pathways. These databases are instrumental in studying secretion and immunoreactivity mechanisms. Before this study, no other research tried to solve misleading localization prediction with the help of patented and immunogenic pathogenic sequences. With the help of patented and immunogenic sequences, PSMS classifies proteins as secreted proteins previously clustered as cytoplasmic. With exploring the patented sequence database, we have observed that for the detection of the pathogens Type 1, Type 2, and Type 4 secretion systems and as a drug target or vaccine studies, effector protein releasing secretion systems (Type 3, Type 5, and outer membrane vesicle proteins) are used. We could not detect immunoreactive proteins in our T6SS database. Although human pathogens, including *Burkholderia mallei* and *B. pseudomallei, Vibrio cholera, Aeromonas hydrophila*, and *Pseudomonas aeruginosa* have a T6SS system in their genome, the recent findings reveal that T6SS is used to target other bacteria and predators, efficiently killing or inhibiting competitors in natural environments [28, 29, and 30]. **We have also realized that EF-TU, Chaperonins, and ATP synthase might involve an unknown vesicular secretion type system for intracellular pathogens. Their vesicular secretions may contain cytoplasmic effector proteins and nucleic acids. While data mining of PUB-MED for the secretion systems of the AntigenDB and USPTO proteins, we have seen this tendency for intracellular pathogens. Identifying this secretion system would improve vaccination strategies for various intracellular pathogens**.

### EF-TU

Although the house-keeping protein EF-Tu proteins are universally known as cytoplasmic proteins, there are various examples of immunoproteomic studies which reported the existence of this protein in secretome and outer membrane vesicles (OMVs), the immunoreactive nature of this protein, and possibilities of its usage as a novel immunogen against bacterial infections) [14, 15, 16 and 17]. The protein must be directed to the periplasm for packaging inside OMV for secretion. Not only proteins (periplasmic and outer membrane proteins) but also lipids, certain metabolites, and even DNA&RNA can be packed heterogeneously inside OMV [18 and 19]. There are some other examples of cytoplasmic proteins leaking into OMVs. As speculated by the authors, the concentration of housekeeping protein inside the cell might be high enough to account for this leak [19 and 20].

Moreover, Caldas et al. [21] showed that E. coli EF-Tu promotes functional folding and protein renaturation after stress, acting as molecular chaperones. Suppose EF-Tu is directed to periplasm for packing inside OMV. In that case, it needs a particular secretion system that might be recognized by PSMS with a T1SS pattern for obligate intracellular pathogens and recognized as an unknown secretion type for facultative intracellular pathogens.

### Chaperonins

Other cytoplasmic proteins, Hsp60 and Hsp10, are immunogenic for various bacterial and fungal pathogens. Various chaperonin studies show that Hsp60 induces a Th-1 immune response and helps pathogens cope with oxidative stress conditions in phagosomes [22]. Also, many researchers have shown the chaperonin vaccine’s protectivity [23, 24, 25]. PSMS patterns show Hsp10 as type 1 secretion associated proteins, where Hsp60 clustered as unknown secretion type associated. Although chaperonins are preventive vaccine candidates, they should be applied as subunit vaccines as they might induce autoimmune cross-reaction with human Hsp60. The chlamydial [26] and helicobacter [27] vaccine preparations showed autoimmune cross-reactions.

### FUTURE WORK

In the future, we are going to use successful PSORT motifs. The PSORT&PSMS hybrid version of the program will be working on a server with suitable immunomodulation future added. We want to focus on toll-like receptor inducing virulence factors as we have realized several pattern recognitions of both DNA&RNA leading to an impaired immune response. The broad-host-range viral structures and the bacterial pathogen exosomes are similar in many ways where a balanced immune response is required to ease the infection. Co-evolution of the pathogens trains pathogens to mutate towards a toll-like response of the host. Our further Analysis will be a toll-like vaccine or therapeutic predictor tool.

## Supporting information

RegEx for Secretory Patterns

## Notes

### Competing Interest Statement

The authors have declared no competing interest.

